# Necroptosis increases with age in the brain and contributes to age-related neuroinflammation

**DOI:** 10.1101/2021.07.23.453580

**Authors:** Nidheesh Thadathil, Evan H Nicklas, Sabira Mohammed, Tommy L Lewis, Arlan Richardson, Sathyaseelan S. Deepa

**Author notes:** **<u>Corresponding Author</u>**: Sathyaseelan S Deepa, Ph.D. Department of Biochemistry and Molecular Biology, Stephenson Cancer Center, Oklahoma Center for Geroscience & Brain Aging, University of Oklahoma Health Sciences Center, 975 NE 10th Street, BRC-1368B, Oklahoma City, OK 73104, USA, (Extn:48393).

## Abstract

Chronic inflammation of the central nervous system (CNS), termed neuroinflammation, is a hallmark of aging and a proposed mediator of cognitive decline associated with aging. Neuroinflammation is characterized by the persistent activation of microglia, the innate immune cells of the CNS, with damage-associated molecular patterns (DAMPs) being one of the well-known activators of microglia. Because necroptosis is a cell death pathway that induce inflammation through the release of DAMPs, we hypothesized that an age-associated increase in necroptosis contributes to increased neuroinflammation with age. The marker of necroptosis, phosphorylated form of MLKL (P-MLKL), and kinases in the necroptosis pathway (RIPK1, RIPK3, and MLKL) showed a region-specific increase in the brain with age, specifically in the cortex layer V and the CA3 region of the hippocampus of mice. Similarly, MLKL-oligomers, which causes membrane binding and permeabilization were significantly increased in the cortex and hippocampus of old mice relative to young mice. Nearly 70 to 80% of P-MLKL immunoreactivity was localized to neurons and less than 10% was localized to microglia, whereas no P-MLKL was detected in astrocytes. P-MLKL expression in neurons was detected in the soma, not in the processes. Blocking necroptosis using *Mlkl*^*-/-*^ mice reduced markers (Iba-1 and GFAP) of neuroinflammation in the brains of old mice and short-term treatment with the necroptosis inhibitor, necrostatin-1s, reduced expression of proinflammatory cytokines, IL-6 and IL-1β, in the hippocampus of old mice. Thus, our data demonstrate for the first time that brain necroptosis increases with age and contributes to age-related neuroinflammation in mice.

## Introduction

Aging is the primary risk factor for the development and progression of various neurodegenerative diseases, including Alzheimer’s disease and Parkinson’s disease [1]. Inflammaging (chronic sterile inflammation) is a characteristic feature of aging and is termed neuroinflammation when it occurs in the central nervous system (CNS). A growing body of preclinical and clinical research have shown that neuroinflammation is a potential mediator of neurodegenerative diseases [2–4]. Neuroinflammation is characterized by the persistent activation of microglia, the innate immune cells of the CNS, and the subsequent sustained release of proinflammatory mediators such as tumor necrosis factor alpha (TNFα), interleukin-6 (IL-6), IL-1β, prostaglandins, reactive oxygen species, and reactive nitrogen species [5]. ‘Resting’ state microglia become activated following exposure to various factors, including damage-associated molecular patterns (DAMPs), which are one of the well-known activators of microglia under sterile conditions [6]. DAMPs bind to pattern recognition receptors (PRRs) such as toll like receptors (TLRs) on innate immune cells in the central nervous system (e.g., microglia), resulting in the induction and secretion of pro-inflammatory cytokines [7,8]. High mobility group box 1 (HMGB1) is one such DAMP that has been shown to facilitate neuroinflammation in aged CNS, and Fonken LK et al. [9] reported that blocking the action of HMGB1 desensitized aged microglia to immune challenges resulting in reduced neuroinflammatory responses following microglia stimulation. HMGB1 is also a known activator of microglia in Alzheimer’s disease and Parkinson’s disease [10].

Necroptosis is a programmed cell death pathway and is a major source of DAMPs [11,12]. Necroptosis is initiated when necroptotic stimuli (e.g. TNF-α, oxidative stress, mTOR/Akt activation etc.) sequentially phosphorylate and activate receptor-interacting protein kinase 1 (RIPK1) and RIPK3, which in turn phosphorylates the pseudokinase mixed lineage kinase domain-like (MLKL) protein [11–13]. Phosphorylation of MLKL leads to its oligomerization, and the oligomerized MLKL then binds to the cell membrane causing its permeabilization and release of cellular components including the DAMPs, which initiate and exacerbate the inflammatory process [12,14]. Necroptosis has been reported to play a role in neuroinflammation in various neurodegenerative diseases such as Alzheimer’s disease [15–17], Parkinson’s disease [18–20], multiple sclerosis [21], and amyotrophic lateral sclerosis [22], as well as brain ischemia [23]. However, it is not known if necroptosis plays a role in age-associated neuroinflammation. Previously, we reported that necroptosis increases with age in the white adipose tissue of mice and interventions that delay aging (dietary restriction or Ames dwarf mice), reduced necroptosis and the expression of pro-inflammatory cytokines [24,25]. We also found that necroptosis was increased in a mouse model of accelerated aging (mice deficient in the antioxidant enzyme, Cu/Zn superoxide dismutase, *Sod1*^*-/-*^ mice) and blocking necroptosis reduced inflammation in the liver of *Sod1*^*-/-*^ mice [26]. Based on these data, we hypothesized that an age-associated increase in necroptosis contributes to increased neuroinflammation with age.

To test our hypothesis, expression of the necroptosis marker, phospho-MLKL (P-MLKL), was assessed in the brains of young and old wild type (WT) mice. We found that expression of P-MLKL increases with age in a region-specific manner in the brain, i.e. layer V of cortex and cornu ammonis 3 (CA3) region of the hippocampus. Nearly 70-80% of P-MLKL immunoreactivity is localized to neurons and less than 10% is localized to microglia, whereas no P-MLKL was detected in astrocytes. Blocking (using *Mlkl*^*-/-*^ *mice)* or inhibiting (using necroptosis inhibitor, necrostatin-1s, Nec-1s) necroptosis resulted in a significant reduction in neuroinflammation in old mice. Thus, our study supports a role of necroptosis in age-associated neuroinflammation.

## Methods

### Animals

C57BL/6JN male wild-type (WT) mice of different age groups (7-, and 22-24-month-old) were received from National Institute on Aging and were maintained at the University of Oklahoma Health Sciences Center (OUHSC) animal care facility for 2 weeks prior to euthanasia. *Ripk3*^*-/-*^ mice (C57BL/6N background) were generated by Vishva M. Dixit (Genentech, San Francisco, CA [27]) and *Mlkl*^*-/-*^ mice (C57BL/6J background) were generated by Warren Alexander (The Walter and Eliza Hall Institute of Medical Research, Australia [28]). We are maintaining colonies of *Ripk3*^*-/-*^ and *Mlkl*^*-/-*^ mice at the OUHSC animal care facility and old *Ripk3*^*-/-*^ and *Mlkl*^*-/-*^ mice for the study were obtained from our colonies. All mice experiments were performed in accordance with the National Institutes of Health’s guidelines and approved by the Institutional Animal Care and Use Committee at OUHSC.

### Immunofluorescence Staining

Mice were anesthetized with isoflurane using anesthesia machine (Kent Scientific Somnosuite) at flow rate of 500 mL/hour for induction and 50 mL/hour for maintenance. Anesthetized mice were perfused with phosphate buffered saline (PBS, pH 7.4) followed by 4% paraformaldehyde (PFA) in PBS and the brain was carefully separated and transferred to 4% PFA for complete fixation for 12 hours followed by 30% sucrose (w/v) solution at 4 °C for 48 hours. The immunofluorescence staining was performed as described by Thadathil et al. [29]. Briefly, 30 µm serial coronal or sagittal sections of brain was sectioned on a cryostat (Leica). Sections were rinsed two times with PBS (pH 7.4) followed by permeabilization with 0.2% triton X-100 for 10 minutes. After washing with PBS, sections were blocked with 3% bovine serum albumin containing 5% donkey or goat serum (Abcam) for 1 hour at room temperature. Sections were incubated overnight at 4 °C with primary antibodies prepared in blocking buffer against phospho-MLKL (Ser345) (Abcam, Cat. No. ab196436), MLKL (Millipore, Cat. No. MABC604), RIPK3 (Novus, Cat. No. NBP1-77299), RIPK1 (Novus, Cat. No. NBP1-77077), neuronal nuclear antigen (NeuN, Abcam, Cat. No. ab104224), ionized calcium binding adaptor molecule 1 (Iba-1, Abcam, Cat. No. ab48004) or glial fibrillary acidic protein (GFAP, Cell Signaling, Cat. No. 3670). After washing the sections with PBS five times, fluorescent labelled secondary antibodies in blocking buffer were added, either alone or in combination, based on the primary antibody and incubated overnight at 4 °C. Following secondary antibodies were used: Goat anti-rabbit Alexa Fluor 647 (Thermofisher Scientific, Cat. No. A-21244), Goat anti-mouse Alexa Fluor 514 (Thermofisher Scientific, Cat. No. A-31555), Donkey anti-goat Alexa Fluor 488 (Thermofisher Scientific, Cat. No. A-11055), and Donkey anti-rat Alexa Fluor 594 (Abcam, Cat. No. ab150156). Stained sections were then washed with PBS and mounted using ProLong™ Diamond antifade mountant (Thermofisher scientific, P36961) which provided labeling of all cell nuclei with DAPI. For the co-localization of P-MLKL with brain cell types (neuron, microglia and astrocytes), serial coronal sections were taken and the number of cells positive for the different antibodies (NeuN, Iba-1 and GFAP) was counted in similar areas (CA3 hippocampus and Layer V cortex) in consecutive serial sections, and values were expressed as the percentage of neurons, astrocytes, and microglia positive to P-MLKL. All the imaging was acquired with a confocal microscope (Zeiss LSM 710) at 200X and 630X magnifications in five non-overlapping fields per mouse and then averaged by an investigator blinded to the groups. Four-six mice per age group was used each staining and images were quantified using ImageJ software (U.S. National Institutes of Health, Bethesda, MD, USA) and normalized with DAPI intensity.

### Quantitative real-time PCR

Relative mRNA levels of necroptosis and inflammatory markers in mouse hippocampus, and cortical region of non-perfused brain was analyzed as described by Mohammed et al. [26]. Briefly, total RNA was extracted using the RNeasy kit (Qiagen, Valencia, CA, USA) from 7-10 mg frozen cortex or hippocampus. First-strand cDNA was synthesized using a high-capacity cDNA reverse transcription kit [ThermoFisher Scientific (Applied Biosystems), Waltham, MA] and quantitative real-time PCR was performed with ABI Prism using Power SYBR Green PCR Master Mix [ThermoFisher Scientific (Applied Biosystems), Waltham, MA]. β-actin was used as the endogenous control. The 2^−ΔΔCT^ comparative method was used to analyze the relative changes in gene expression and represented as fold change. The sequence of primers used for the study are: TNF-α (forward 5’-CACAGAAAGCATGATCCGCGACGT-3’; reverse 5’-CGGCAGAGAGGAGGTTGACTTTCT-3’), IL-6 (forward 5’-TGGTACTCCAGAAGACCAGAGG-3’, reverse 5’-AACGATGATGCACTTGCAGA-3’), IL-1β (forward 5’-AGGTCAAAGGTTTGGAAGCA-3’, reverse 5’-TGAAGCAGCTATGGCAACTG-3’), and β-actin (forward 5′-ATGGATGACGATATCGCTG-3′, reverse 5′-GTTGGTAACAATGCCATGTTC-3′).

### Western Blotting

Cortex and hippocampus were collected from non-perfused brain collected during sacrifice and were immediately frozen in liquid nitrogen and stored at −80 °C until use. For western blotting, 15 mg tissues were homogenized in RIPA lysis buffer (ThermoFisher Scientific, Waltham, MA) containing phosphatase and protease inhibitor cocktail (Goldbio Cat. No. GB-108-2). Western blotting was performed using 40 μg protein/well as previously described (Mohammed et al. 2021). Primary antibodies against the following proteins were used: Phospho-MLKL (Ser345) (Abcam, Cat. No. ab196436); MLKL (Millipore, Cat. No. MABC604), β-tubulin (Cat. No. T5201) and GAPDH (Cat. No. G8795) were purchased from Sigma-Aldrich (St. Louis, MO). HRP-linked anti-rabbit IgG, HRP-linked anti-mouse IgG or HRP-linked anti-rat IgG were used as secondary antibodies (Cell Signaling Technology, Danvers, MA). Membranes were imaged with ChemiDoc imaging system (Bio-Rad) and quantified using ImageJ software (U.S. National Institutes of Health, Bethesda, MD, USA).

### MLKL-oligomers detection by western blotting

MLKL oligomerization in hippocampus and cortex was detected by western blotting under non-reducing conditions following the protocol described by Cai and Liu [30]. Hippocampus and cortex were homogenized in RIPA lysis buffer and protein quantification was performed as described above. Forty micrograms of total hippocampus and cortex protein was mixed with 2X Laemmli buffer (Bio-Rad) without β-mercaptoethanol and boiled at 95 °C for 5 minutes. The samples were then loaded into 10% SDS-PAGE gel and western blotting was performed as described above. Anti-MLKL antibody (Millipore, Cat No. MABC604) was used to detect MLKL-oligomers.

### Nec-1s treatment

Three groups of mice were used for the study: young WT mice (7-months) treated with vehicle (n=8), old WT mice treated with vehicle (n=6) and old WT mice treated with Nec-1s (n=8). On the first day of treatment, mice were given a single intraperitoneal injection of 10 mg/kg Nec-1s (7-Cl-O-Nec-1; Focus Biomolecules, Plymouth Meeting, PA) or vehicle, followed by administration of Nec-1s in drinking water for 30 days [21,26]. For supplementation in drinking water, Nec-1s was first dissolved in dimethylsufoxide (DMSO, 50% w/v) and was transferred into 35% polyethylene glycol (PEG) solution and was suspended in water containing 2% sucrose (final concentration: 0.5 mg/mL of Nec-1s). Daily consumption of Nec-1s based on this protocol is reported to be 2.5–5 mg/day [22]. After 30 days, cortex and hippocampus and were collected from non-perfused brain during sacrifice and were immediately frozen in liquid nitrogen and stored at -80 °C until use.

### Statistical analyses

Ordinary one-way ANOVA with Tukey’s post hoc test and Student’s t test was used to analyze data as indicated. P< 0.05 is considered statistically significant.

## Results

### Necroptosis markers increase with age in a region-specific manner in the brain

To measure the changes in necroptosis with age in the brain, we first measured the expression of P-MLKL, which is involved in the final step in necroptosis [31,32], in the brains of young (7-month-old) and old (22 to 24-month-old) WT mice. As shown in **Figure 1A**, expression of P-MLKL was increased in the brains of old WT mice relative to young WT mice as measured by immunofluorescence staining. Importantly, P-MLKL showed a region-specific expression in the brains of old WT mice: i.e. P-MLKL expression was observed primarily in layer V of the cortex (**Figure 1B**) and in the CA3 region of the hippocampus (**Figure 1C**). Due to region-specific expression of P-MLKL, we focused our study on the cortex layer V and the CA3 region of the hippocampus for further immunofluorescence analyses. Quantification of percentage of cells that express P-MLKL showed that P-MLKL expression was 5-fold higher in cortex layer V and 4-fold higher in the CA3 region of the hippocampus of old WT mice relative to young WT mice (**Figure 2A**).

**Figure 1.**
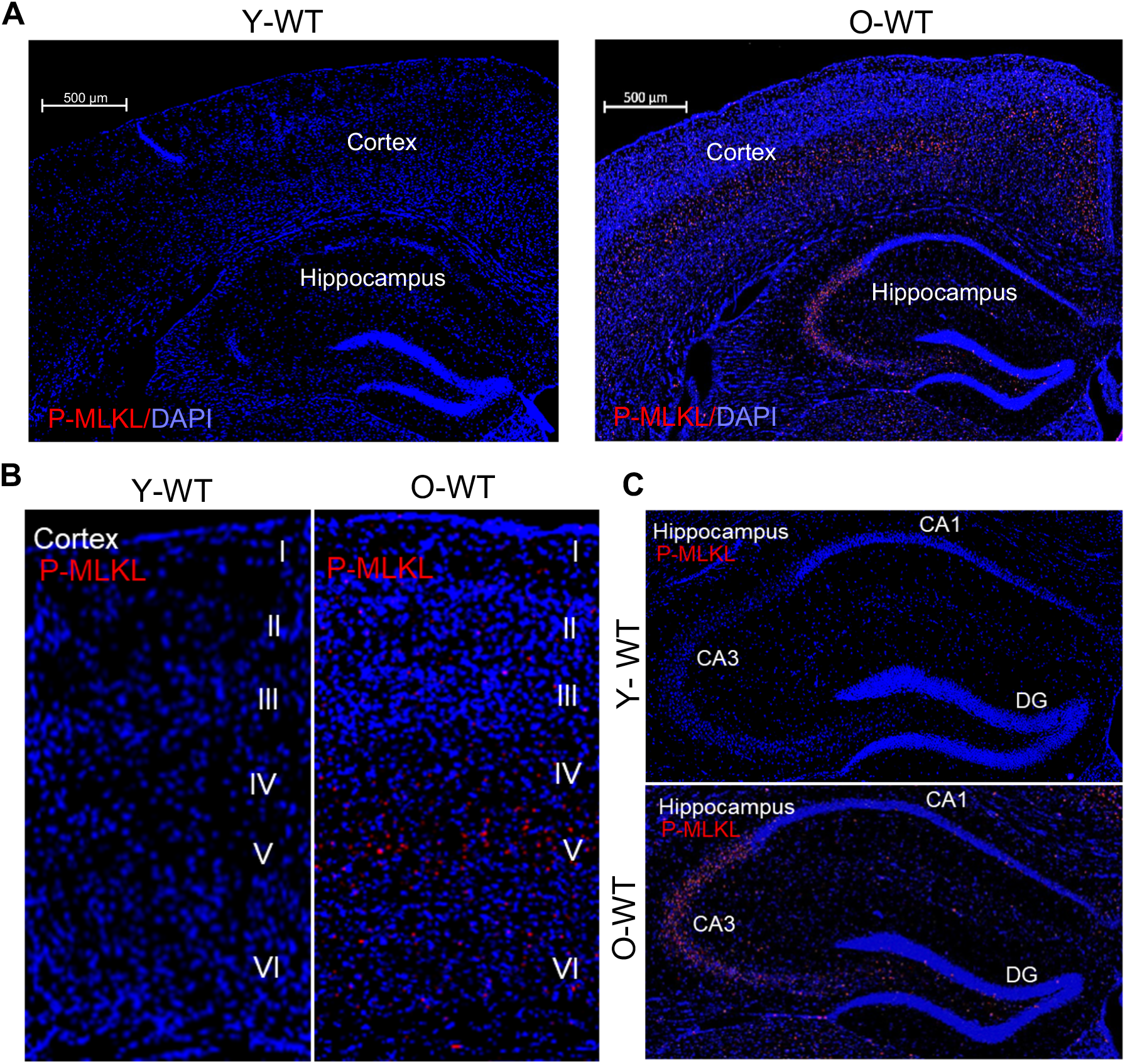
Expression of P-MLKL in different brain regions of young and old wild type mice. **(A)** Confocal microphotographs showing expression of P-MLKL in the brains of young wild type (WT) and old WT mice. P-MLKL staining is shown in red and DAPI staining in blue. Scale bar: 500 μm. Enlarged images showing P-MLKL expression in the cortex **(B)** and the hippocampus **(C)** of young and old WT mice.

**Figure 2.**
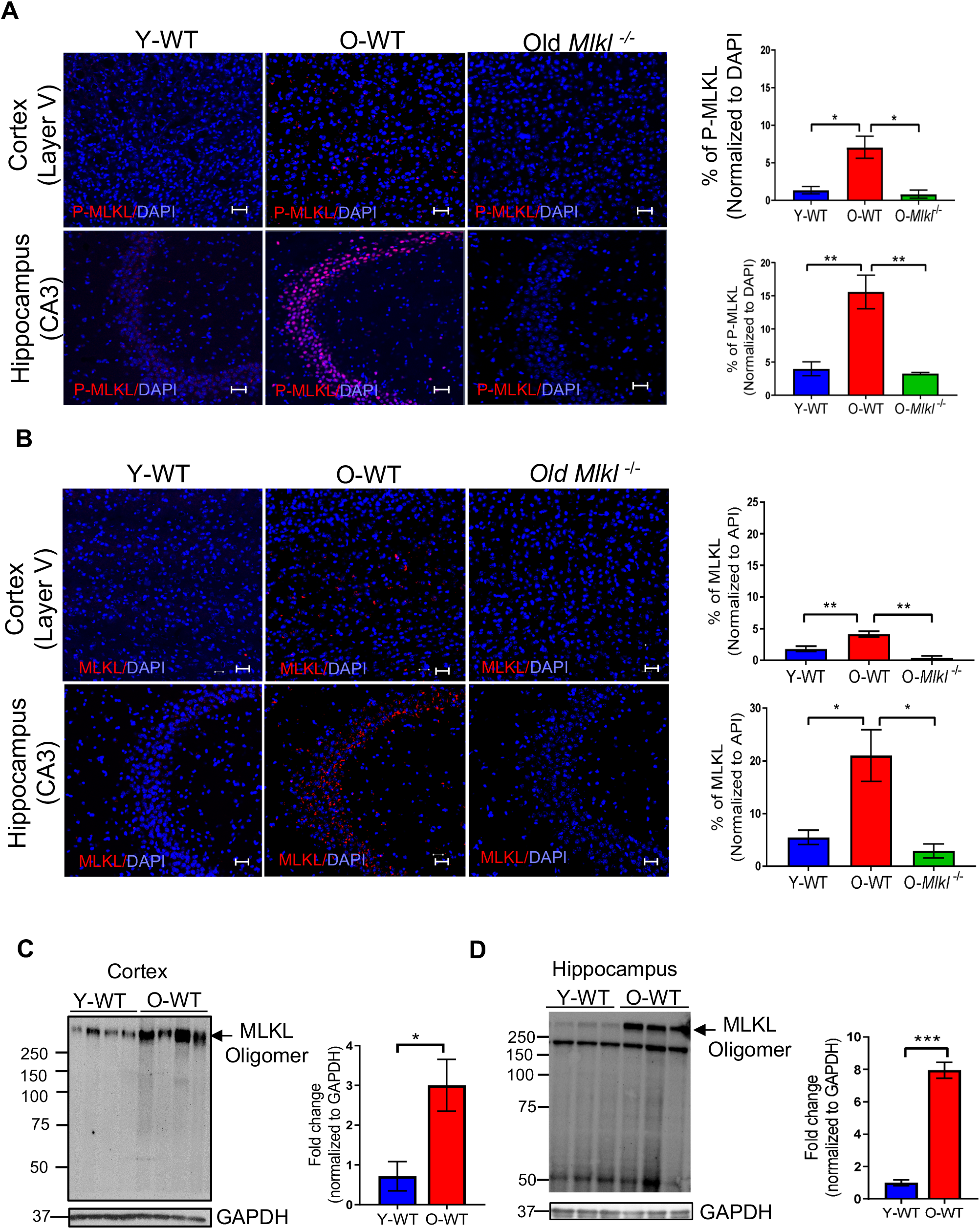
Changes in the expression of P-MLKL, MLKL, and MLKL-oligomers in the brain with age. Confocal imaging showing expression of P-MLKL **(A)** and MLKL **(B)** in the layer V of the cortex and the CA3 region of the hippocampus of young WT (Y-WT), old WT (O-WT), and old *Mlkl*^*-/-*^ mice. P-MLKL and MLKL staining is in red and DAPI staining is in blue (*left panel)*. Graphical representation of the percentage of P-MLKL or MLKL normalized to DAPI Y-WT (blue bar), O-WT (red bar), and old *Mlkl*^*-/-*^ mice (green bar) are shown in the *right panel*. Data were obtained from 7 to 8 mice per group and are expressed as the mean ± SEM. (ANOVA, * P ≤ 0.05). Scale bar: 20 μm. Western blot analysis of MLKL-oligomers in Y-WT and O-WT cortex **(C)** and hippocampus **(D)**. Quantification of MLKL-oligomers normalized to GAPDH is represented on the right side of representative blots. Data were obtained from 5 to 7 mice per group and are expressed as the mean ± SEM. * p ≤ 0.05, **p≤ 0.01, ***p≤ 0.001.

To test the specificity of P-MLKL antibody, brain sections from old *Mlkl*^*-/-*^ mice were used as controls. As shown in **Figure 2A** (3^rd^ panel), we did not observe any staining for P-MLKL in the brain sections from old *Mlkl*^*-/-*^ mice, demonstrating that the positive staining for P-MLKL in old WT brains was specific for P-MLKL. Similar to P-MLKL, expression of MLKL was significantly upregulated in the cortex layer V (2-fold) and the CA3 region of the hippocampus (4-fold) of old WT mice relative to young WT mice, whereas MLKL expression was absent in the brains of old *Mlkl*^*-/-*^ mice (**Figure 2B**). Analysis of MLKL transcript levels showed that levels of MLKL were significantly higher in the hippocampus (1.5-fold) of old WT mice relative to young WT mice, suggesting that increased transcription might contribute to increased MLKL expression. Even though *Mlkl* transcript levels were increased in the cortex of old WT mice relative to young WT mice, the difference was not statistically significant (**Supplementary Figure S1A**).

Phosphorylation of MLKL leads to its oligomerization, which results in the translocation of MLKL-oligomers to the cell membrane causing membrane permeabilization and release of DAMPs [32]. Therefore, we measured the levels of MLKL-oligomers in the cortex and hippocampus of young and old WT mice. Consistent with the age-associated increase in the expression of P-MLKL, MLKL-oligomers showed a significant increase in the cortex (3-fold) and hippocampus (8-fold) of old WT mice relative to young WT mice (**Figure 2C-D**), providing additional evidence for increased necroptosis with age in these brain regions.

We also assessed expression of RIPK3 and RIPK1, which form necrosome complex once phosphorylated and facilitate MLKL phosphorylation [33,34]. Expression of RIPK3 and RIPK1 were significantly upregulated in cortex layer V (2.5- and 10-fold) and the CA3 region of the hippocampus (3.5- and 13.7-fold) of old WT mice relative to young WT mice similar to what we observed for MLKL (**Supplementary Figures S2A and S2B**). Because of the non-specific binding of anti-P-RIPK3 and anti-P-RIPK1 antibodies, we were unable to assess the expression of RIPK3 and RIPK1 phosphorylation in the cortex layer V and CA3 region of the hippocampus. Transcript levels of *Ripk3* and *Ripk1* were similar in the cortex of young and old WT mice (**Supplementary Figure S1B**). In the hippocampus of old mice, the level of *Ripk1* transcript was significantly higher relative to young WT mice (1.2 fold), whereas *Ripk3* transcript levels were similar in young and old WT mice (**Supplementary Figures S1C**). Thus, aging is associated with increased expression of RIPK1, RIPK3, and MLKL and the activation of MLKL (phosphorylation and oligomerization) in the cortex layer V and the CA3 region of the hippocampus of mice.

### Necroptosis markers in the cortex and hippocampus are mainly localized to neurons

To identify the cell types that express necroptosis marker, we performed co-immunostaining of P-MLKL with markers of neurons (neuronal nuclear antigen, NeuN, [35]), microglia (ionized calcium binding adaptor molecule 1, Iba-1, [36]), and astrocytes (glial fibrillary acidic protein, GFAP, [37]). As shown in **Figure 3A**, P-MLKL staining colocalized with NeuN (top panel) and Iba-1 (middle panel) in the cortex layer V of old WT mice. No colocalization of P-MLKL staining was observed with GFAP (bottom panel). Quantification of the percentage of P-MLKL positive cells in the cortex showed that 72% of P-MLKL was localized to neurons, 9% to microglia, and no P-MLKL was colocalized to astrocytes. In the CA3 region of the hippocampus, 79% of total P-MLKL staining was localized to neurons, 4% was localized to microglia, and no P-MLKL expression was detected in astrocytes (**Figure 3B**). Co-immunostaining of MLKL, RIPK3 or RIPK1 with NeuN showed that these three proteins are localized to the neurons in old WT mice **(Supplementary Figure S3)**. Thus, in the cortex layer V and the CA3 region of the hippocampus of old WT mice, neurons were identified as the major cell type that express increased necroptosis markers.

**Figure 3.**
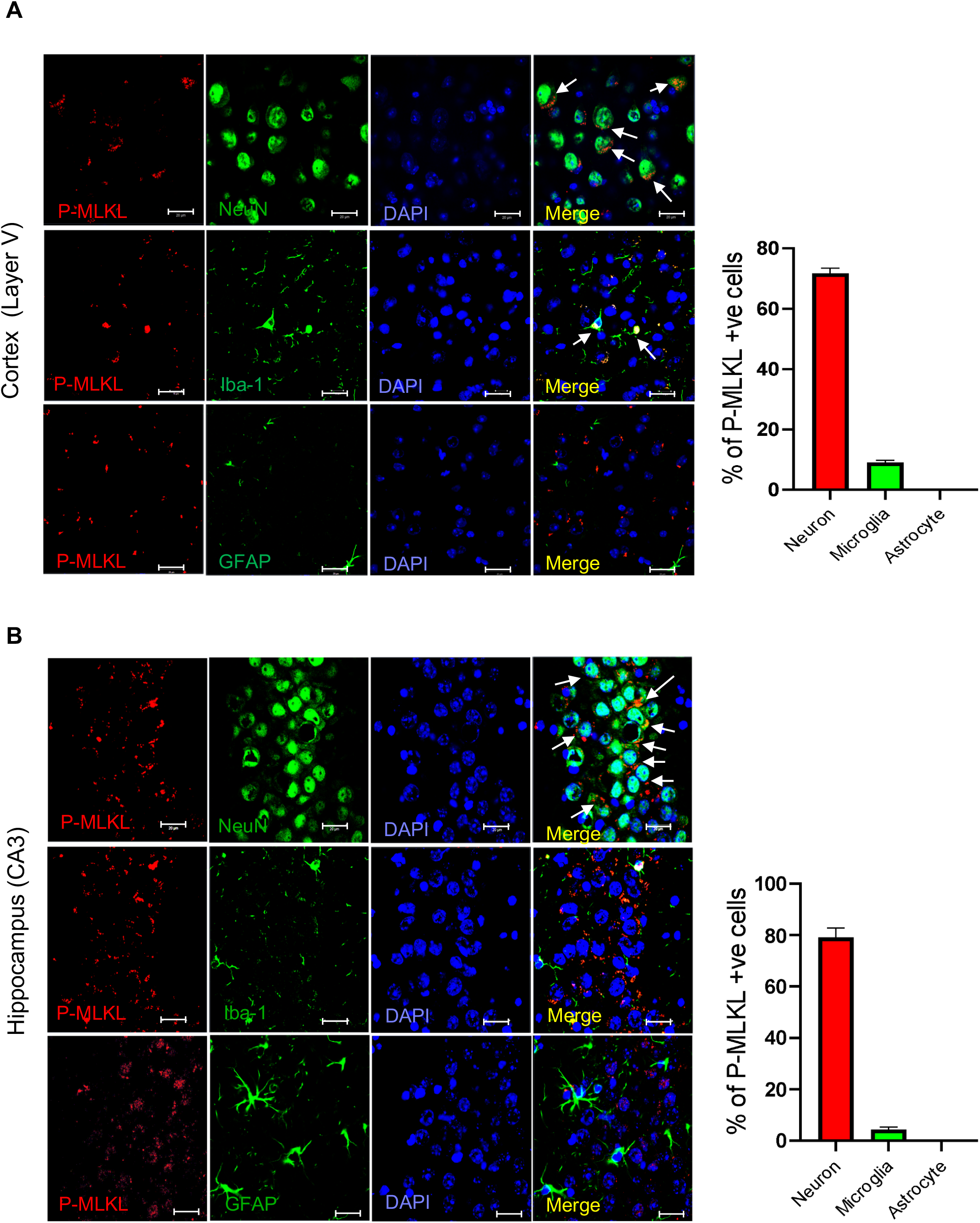
Co-localization of P-MLKL in various brain cell types in old WT brain. Confocal images showing expression of P-MLKL in neurons, microglia and astrocytes in the cortex layer V **(A)** and CA3 region of the hippocampus **(B)**. Double immunofluorescence staining and confocal micrographs are shown for neurons (NeuN, green, top panel), microglia (Iba-1, green, middle panel), and astrocytes (GFAP, green, bottom panel) with P-MLKL (red) and DAPI (blue). White arrows indicate the co-localization of P-MLKL with the various cell types. Graphical representation of percentage of P-MLKL positive cells co-localized with brain cell type markers (right panel). Scale bar :20 μm. Data were obtained from 5 to 7 mice and are expressed as the mean ± SEM. * p ≤ 0.05.

Next, we assessed localization of P-MLKL in the neurons of old WT mice using phase contrast microscopy. Co-immunostaining of P-MLKL with NeuN and DAPI showed that P-MLKL was localized to the soma of neurons in the cortex layer V of old WT mice (**Figure 4A**). Because the NeuN antibody predominantly stains the nuclei of post-mitotic neurons [38], we confirmed the localization of P-MLKL in the neurons by the co-localization of P-MLKL with the nuclear stain, DAPI. As shown in **Figure 4B**, P-MLKL was mainly localized outside the nuclei. Similarly, in the CA3 region of the hippocampus, P-MLKL was localized to the soma region of neurons, outside the nucleus (**Figures 4C and 4D**). Thus, aging is associated with increased expression of P-MLKL, which is found mainly in the soma of neurons.

**Figure 4.**
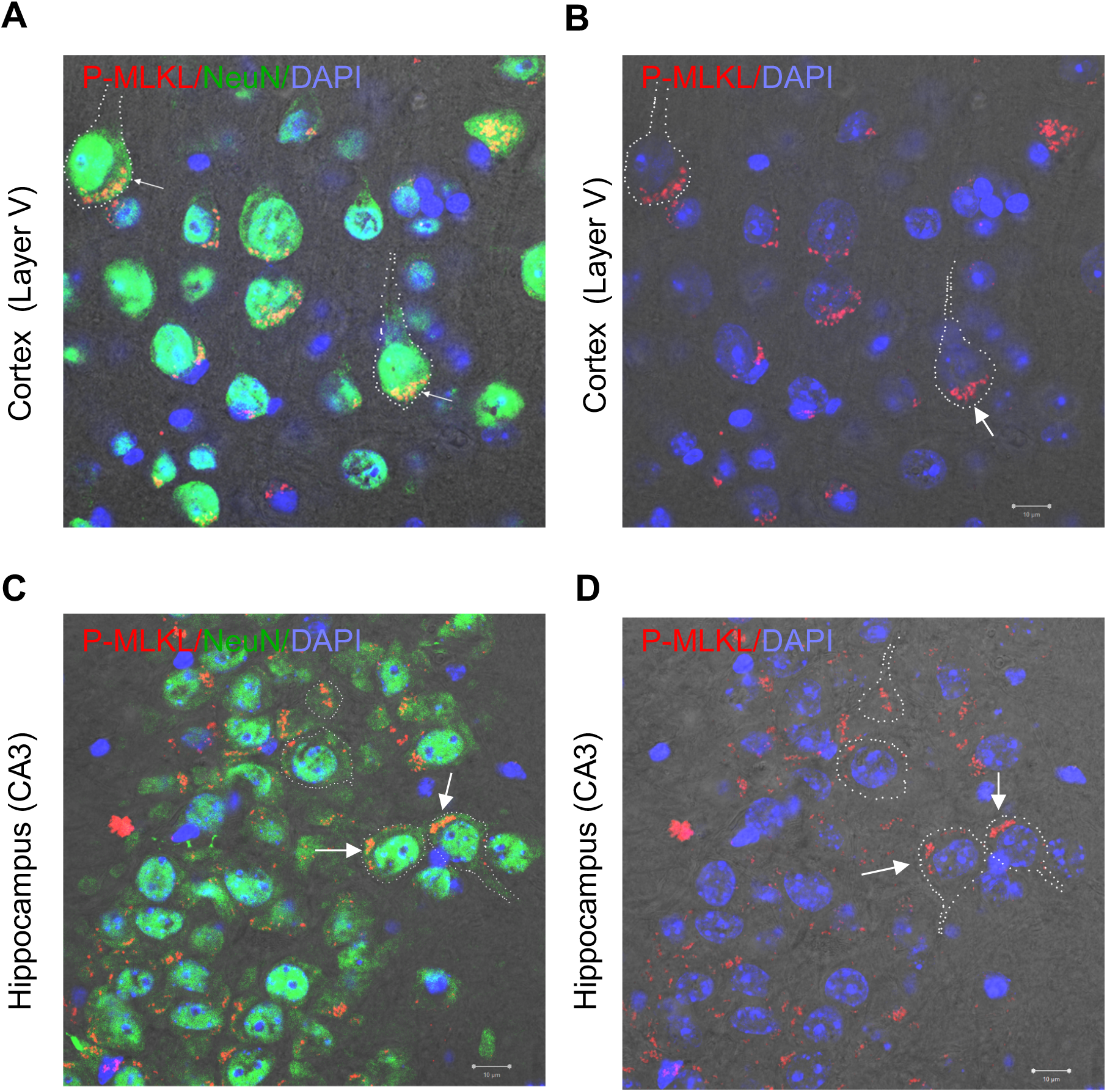
Cellular localization of P-MLKL in neurons. T-PMT (photomultiplier for transmitted light) micrographs showing P-MLKL in red and DAPI in blue in the layer V of the cortex **(A)** and the CA3 region of the hippocampus **(C). Panels B** and **D** show the double immunofluorescence images of P-MLKL with neuronal marker NeuN in the layer V Cortex and the hippocampal CA3 region, respectively. Dotted line indicates the periphery of the neuron and arrow indicates the location of P-MLKL in the cell. Scale bar :10 μm. Representative micrographs obtained from 4 to 6 mice.

### Effect of necroptosis inhibition on neuroinflammation in old mice

Neuroinflammation is a hallmark of the aging brain and age-related neurogenerative diseases (e.g., Alzheimer’s disease, AD) and has been associated with cognitive decline in these conditions [39,40]. Hippocampus is the major brain region associated with cognition, and chronic inflammation in hippocampus has been shown to be associated with cognitive impairment in mice [41,42]. Because necroptosis has been shown to be a major pathway involved in inflammation [16], we directly tested the role of the age-related increase in necroptosis (i.e., P-MLKL) on neuroinflammation by measuring two widely used markers of neuroinflammation, Iba-1 and GFAP [43,44] in the CA3 region of the hippocampus in young and old WT mice and old *Mlkl*^*-/-*^ mice. Consistent with previous reports, we found that expression of Iba1 [45] and GFAP [46] was increased in the CA3 region of the hippocampus of old WT mice relative to young WT mice (**Figure 5**). Iba-1 and GFAP expression was significantly increased 2-fold and 3.4-fold, respectively in old WT mice relative to young WT mice (**Figures 5A and 5B**). To determine if this increased neuroinflammation was due to increased necroptosis, which we observed in the CA3 region of the hippocampus of the old WT mice, we measured Iba-1 **(Figure 5A)** and GFAP **(Figure 5B)** expression in the brains of old *Mlkl*^*-/-*^ mice, which we showed blocked increased necroptosis (P-MLKL) in the brains of old WT mice (**Figure 2A**). As shown in **Figures 5A and 5B**, the expression of Iba-1 and GFAP was significantly reduced in old *Mlkl*^*-/-*^ mice.

**Figure 5.**
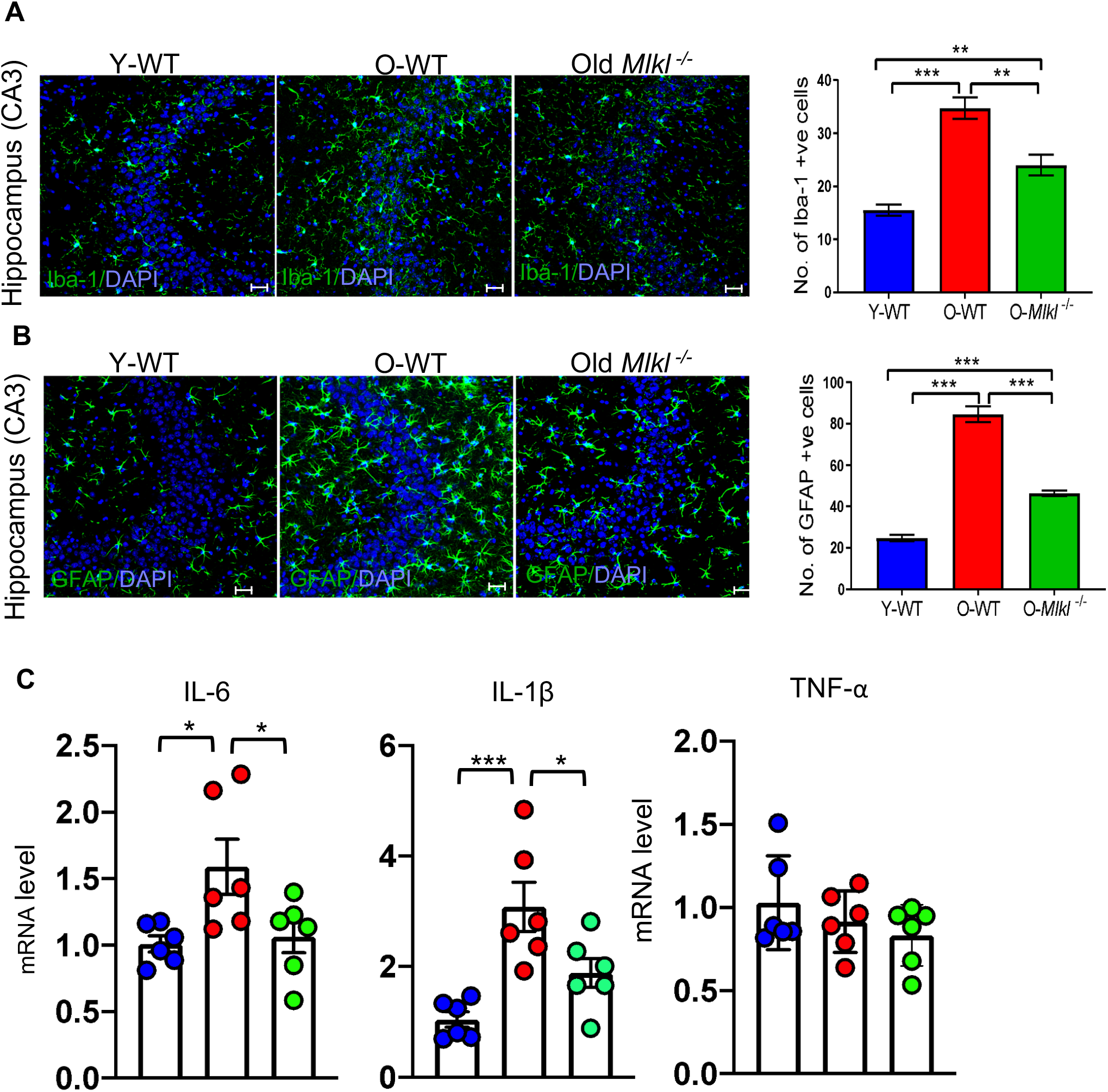
Effect of blocking necroptosis on the expression of neuroinflammatory markers in the brains of old WT mice. Confocal images (left panel) showing the expression of the following: Iba-1 (green) and DAPI (blue) **(A)** and GFAP (green) and DAPI (blue) **(B)** in the hippocampal CA3 region of Y-WT, O-WT, and old *Mlkl*^*-/-*^ mice. Graphical representation of the number of Iba-1 or GFAP positive cells in are shown in the right panel. **(C)** Transcript levels of pro-inflammatory cytokines IL-6, IL-1β and TNF-α in the hippocampus of vehicle treated Y-WT (blue), vehicle treated O-WT (red), and Nec-1s treated O-WT (green) mice. Data were obtained from 5 to 7 mice per group and are expressed as the mean ± SEM. * p ≤ 0.05, **p≤ 0.01, ***p≤ 0.001.

We also tested the effect of inhibiting necroptosis on the expression of proinflammatory cytokines in the hippocampus of old WT mice by treating old WT mice with vehicle or necrostatin-1s (an inhibitor of necroptosis that targets RIPK1) for 1 month. Nec-1s treatment has been shown to significantly reduce neuroinflammation associated with multiple sclerosis, amyotrophic lateral sclerosis and AD [16,21,22]. In these experiments, we measured the transcript levels of TNFα, IL-6, and IL-1β in the hippocampus of old mice treated with Nec-1s because these pro-inflammatory cytokines have been shown to be associated with inflammaging [47–49]. In the hippocampus of old WT mice, transcript levels of IL-6 (1.5-fold) and IL-1β (3-fold) were significantly elevated relative to young WT mice, and Nec-1s treatment resulted in a significant reduction in the transcript levels of IL-6 and IL-1β (**Figure 5C**). However, neither age nor Nec-1s treatment had any effect on TNFα transcript levels (**Figure 5C**). Together, these data indicate that the age-related increase neuroinflammation in the hippocampus of old mice is at least partially due to increased necroptosis.

## Discussion

In contrast to apoptosis, which is a non-inflammatory mode of cell death, necrosis is a highly pro-inflammatory form of cell death due to the release of DAMPs from the dying cells [50,51]. In 2005, Degterev et al. [11] reported that cells are capable of activating programmed necrosis, termed necroptosis, in the absence of intracellular apoptotic signaling. Over the past decade, necroptosis has been reported to play a major role in inflammation in a variety of conditions and diseases, e.g., atherosclerosis, fatty liver disease, macular degeneration and neurogenerative diseases such as Alzheimer’s disease and Parkinson’s disease [25]. In 2018, we reported the first evidence that necroptosis increases with age in the white adipose tissue of mice [24]. Because necroptosis and inflammation were reduced by dietary restriction and in Ames dwarf mice, we suggested that necroptosis plays a role in inflammaging [24,25]. Subsequently, we reported that necroptosis was increased in the liver of a mouse model of accelerated aging, mice deficient in the antioxidant enzyme, *Sod1* [26]. Inhibition of necroptosis reduced expression of proinflammatory cytokines in the livers of *Sod1*^*-/-*^ mice, supporting a role of necroptosis in age-associated inflammation.

The age-related increase in chronic inflammation in the central nervous system, termed neuroinflammation, is recognized to be an important factor in the age-related decline in cognition [52] and is a major risk factor in neurodegenerative diseases, such as Alzheimer’s disease. The goal of this study was to determine if changes in necroptosis occurred in brain with age and to assess the role of necroptosis in neuroinflammation. First, we evaluated the expression of necroptosis marker, P-MLKL, in the brains of young and old mice and found that P-MLKL expression was increased with age in the brain. Consistent with increased P-MLKL, we observed an increase in the levels of MLKL-oligomers in the hippocampus and cortex of old mice. Oligomerization of MLKL is a key determinant of its membrane translocation and membrane permeabilization [32]. In addition, we observed that the expression of RIPK1 and RIPK3, the kinases that form the necrosome complex and recruits MLKL for phosphorylation were also elevated in the brains of old mice. Together, these data strongly support an age-related increase in necroptosis activation in the brain. Increased necroptosis has been reported in the postmortem brains of Alzheimer’s disease patients, i.e. an increase in the expression of RIPK1, RIPK3, MLKL and P-MLKL. An increase in necroptosis (i.e. P-MLKL expression) has also been reported in the brains of patients with Parkinson’s disease and multiple sclerosis [18,53].

Interestingly, we observed that the age-related increase in P-MLKL expression was not evenly distributed throughout the brain but occurred in specific brain regions of old mice, e.g., the CA3 region of the hippocampus and the layer V of the cortex. A region-specific expression of P-MLKL has also been observed in the brain in neurodegenerative diseases. For example, the hippocampus and temporal gyrus were identified as the regions that had the highest expression of P-MLKL in Alzheimer’s disease [15], increased expression of P-MLKL was reported in the cortical layers II-III in multiple sclerosis [53], and P-MLKL expression was found in substantia nigra in Parkinson’s disease [18]. While the reason(s) for region-specific necroptosis in the brain in aging and neurogenerative diseases is not clear, it is possible that increased stress or factors that induce necroptosis are more predominant in these brain regions under disease conditions and aging. TNFα, oxidative stress, and mTOR (mammalian target of rapamycin) activation are known to induce necroptosis and the expression or activation of these factors increases with age and neurodegenerative diseases [25]. TNF-α, a pro-inflammatory cytokine and the master regulator of the innate immune response, is a well-known activator of necroptosis [54]. In humans, an increase in peripheral TNFα has been reported with age and is strongly associated with gray matter loss and cognitive decline [55]. A role of TNF-α in neuronal necroptosis is supported by the report that blocking necroptosis by targeting *Ripk3* (using *Ripk3*^*-/-*^ mice) reduced neuronal loss mediated by the intracerebroventricular injection of TNFα. Importantly, in wild type mice, intracerebroventricular injection of TNFα reduced neuronal density especially in the CA3 region of hippocampus suggesting increased susceptibility of neurons in the CA3 regions of hippocampus to TNFα-mediated cell death [56]. Picon et al [53] showed that in the brains of multiple sclerosis patients, TNF receptor 1 (TNFR1) and P-MLKL are specifically enriched in layer II-III of cortex and P-MLKL colocalized with TNFR1 in neurons in layer II-III, further supporting a role of TNFα signaling in necroptosis activation.

Several studies support a role of oxidative stress in the induction of necroptosis. For example, oxidative stress is shown to induce necroptosis of primary cortical neurons in cell culture studies [57] and increased oxidative stress is associated with dopaminergic neuronal necroptosis in Parkinson’s disease [19]. Oxidative damage has been reported to increase with age to different levels in different brain regions with hippocampus being the brain region most impacted [58,59]. In addition, oxidative stress has been proposed as a mediator of neurodegeneration in several neurogenerative diseases [60]. Within the hippocampus, both CA1 and CA3 regions are reported to have increased oxidative stress in a mouse model of Alzheimer’s disease (*APP*^*NL-G-F*^ mice) or in *Sod2*^*+/-*^ mice [61,62]. Surprisingly, neurons in the CA1 region of hippocampus are highly vulnerable to cell death mediated by oxidative stress, whereas neurons in the CA3 region of hippocampus are resistant to oxidative stress mediated cell death [60,63,64]. Therefore, age-associated increase in oxidative stress might not be a major factor mediating the age-related increase in necroptosis in the CA3 region of the hippocampus. The activation of mTOR is yet another proposed mediator of necroptosis. In hippocampal neuronal cell line HT22, induction of necroptosis was blocked by the combined treatment of Akt and mTOR inhibitors [65]. In a mouse model of cerebral palsy, mTOR inhibitor rapamycin prevented neuroinflammation and neuronal cell death [66]. In Alzheimer’s disease patients and mouse models, hyperactive mTOR signaling has been reported in the brain regions affected by the disease [67,68]. Feeding AD mice with rapamycin, a mTOR inhibitor, has been shown to improve cognition, reduce Aβ and tau pathology, and reduce neuroinflammation (microglia activation) [69,70]. mTOR activity is reported to increase with age in tissues such as liver, muscle, heart, and adipose tissue [71], whereas in the hippocampus, activity of mTOR and mTOR signaling is reported to decline with age [72]. Based on the current literature, we speculate that age-associated increase in TNFα is most likely to be the factor contributing to necroptosis occurring in specific regions of the aging brain. Whether TNFα receptor expression increases in the hippocampal CA3 region or layer V of cortex with age needs to be tested.

To determine what cell types in the CA3 region of the hippocampus and layer V of the cortex showed the increase in necroptosis with age, we measured the co-localization of P-MLKL with markers of neurons (NeuN), microglia (Iba-1), and astrocytes (GAFP) using immunofluorescence. Neurons were identified as the major cell type that express P-MLKL, e.g., almost 80% of the total P-MLKL was localized to neurons. We detected no P-MLKL expression in GFAP-positive astrocytes in the brains of old mice. Expression of RIPK1, RIPK3, and MLKL were also increased in the neurons of old mice suggesting greater levels of necroptosis machinery in neurons with age. Our data are similar to reports of necroptosis in neurodegeneration where neurons were identified as the major cell type that express P-MLKL in the brains of patients with Alzheimer’s disease [15], multiple sclerosis [53], Parkinson’s disease [18], ischemic brain [73,74], and in Japanese encephalitis virus infection [75]. In the brains of patients with multiple sclerosis, microglia do not undergo necroptosis due to the lower expression of MLKL, whereas oligodendrocytes that have a higher expression of MLKL undergo necroptosis [21]. In our study, we found that MLKL expression is higher in neurons of old mice, whereas only very few microglia expressed MLKL. Therefore, it is possible that in neurogenerative diseases as well as in aging, the cell types that have a robust expression of MLKL (as well as RIPK1 and RIPK3) will have high P-MLKL expression and necroptosis. Picon et al [53] reported that only neurons, not microglia or astrocytes, express TNFR1 in the brains of multiple sclerosis patients, again suggesting that increased TNFα-signaling might promote increased necroptosis in neurons relative to other cell types in aging. Whereas neuronal loss is a characteristic feature of neurogenerative diseases, neuronal loss is minimal in brain aging [76]. Therefore, we speculate that an increase in the levels of P-MLKL in the neurons of old mice might cause membrane leakiness, without killing neurons, to release DAMPs. Cell culture studies have shown that in dendritic cells, increased P-MLKL can cause inflammation without promoting cell death. This is believed to occur through the release of extracellular vesicles (EV) containing P-MLKL [74,77,78]. The necroptotic EVs can be phagocytosed by immune cells to increase cytokine and chemokine secretion, which could lead to increased inflammation [79]. Therefore, it is possible that neurons in old mice may avoid cell death by increasing the release of P-MLKL containing EVs and induce neuroinflammation, however, this possibility needs to be tested.

In addition to neurons, a small percentage of microglia also showed increased expression of P-MLKL (less than 10% of the total P-MLKL) in the brains of old mice. In the brains of AD patients, nearly 30% of P-MLKL is colocalized with microglial marker, Iba1 [15]. Microglial necroptosis is reported to cause neuroinflammation and exacerbate neuronal damage in retinal degeneration [80], and retinopathy [81]. In addition, necroptosis of proinflammatory microglia is involved in remyelination of CNS [82]. Whether a subpopulation of microglia undergo necroptosis or microglial necroptosis play a role in neuroinflammation in the aging brain needs to be investigated. Even though astrocytes are reported to undergo necroptosis in a mouse models of spinal cord injury and cerebral ischemia [73,74], we did not see any P-MLKL expression co-localize with GFAP-positive astrocytes from old mice. In the brains of AD patients, nearly 10% of total P-MLKL was observed to colocalize with GFAP-positive astrocytes. Necroptosis has been reported for other cell types in the brain. For example, oligodendrocytes undergo necroptosis in multiple sclerosis [21], whereas necroptosis occurs in vascular endothelial cells in stroke [83]. Whether oligodendrocytes or vascular endothelial cells undergo necroptosis in aging needs to be tested in the future.

Because changes in necroptosis have been shown lead to increased inflammation in various conditions and diseases, we first measured classical markers of neuroinflammation (Iba-1 and GFAP) in the hippocampus of young and old mice. As has been previously reported, these markers were increased in the hippocampus of the old mice [84–87], which correlated with the increase in necroptosis. We also observed an increase in the mRNA transcripts for IL-6 And IL-1β in the hippocampus of old mice. To determine if necroptosis was responsible for neuroinflammation in old mice, we blocked (using *Mlkl*^*-/-*^ *mice*) or inhibited (using necrostatin-1s) necroptosis in old mice. We found that blocking or reducing necroptosis reduced markers of neuroinflammation in the brains of old mice supporting a role of necroptosis in neuroinflammation associated with age. Blocking necroptosis by targeting *Mlkl* has been shown by Bian et al. [75] to reduce neuroinflammation in response to Japanese encephalitis virus infection. Similarly, inhibition of necroptosis using Nec-1s, an allosteric inhibitor of RIPK1, has been shown to reduce neuroinflammation in Alzheimer’s disease, Parkinson’s disease, multiple sclerosis, and amyotrophic lateral sclerosis [15,18,21,22]. However, future studies targeting Mlkl specifically in neurons will be required to address the role of neuronal necroptosis in age-related neuroinflammation.

In summary, our study demonstrates for the first time that markers of necroptosis are increased in specific regions of the aging brain and shows that necroptosis plays a role in neuroinflammation associated with aging. Our finding that a short term Nec-1s treatment late in life can effectively block neuroinflammation suggests that blocking necroptosis would be an effective strategy to reduce age-associated neuroinflammation and possibly retard cognitive decline in aging. Nec-1s is extensively used in preclinical studies and several other RIPK1 inhibitors (e.g. DNL474, DNL758) are currently in clinical trials for neurodegenerative diseases suggesting the translational potential of these inhibitors for age-related neuroinflammation and cognitive decline [88].

## Acknowledgments

The authors would like to thank Imaging core facility at the Oklahoma Medical Research Foundation for confocal imaging services.

## Declarations

### Availability of data and material

Available upon request

### Code availability

Not applicable

### Authors’ contributions

N.T. performed the experiments, analyzed data, and prepared figures; E.N. performed real-time qPCR experiments and assisted with western blotting, S.M. edited the manuscript; T.L.L.Jr. gave advise for immunofluorescence staining, helped with data interpretation and edited the manuscript; A.R. gave critical comments and suggestions for the manuscript and edited the manuscript; and S.S.D designed the experiments, wrote and edited the manuscript.

### Ethics approval

Not applicable

### Consent to participate

Not applicable

### Consent for publication

All authors have agreed to the content of this manuscript for publication.

## Supplementary Figure Legends

**Supplementary Figure 1:**
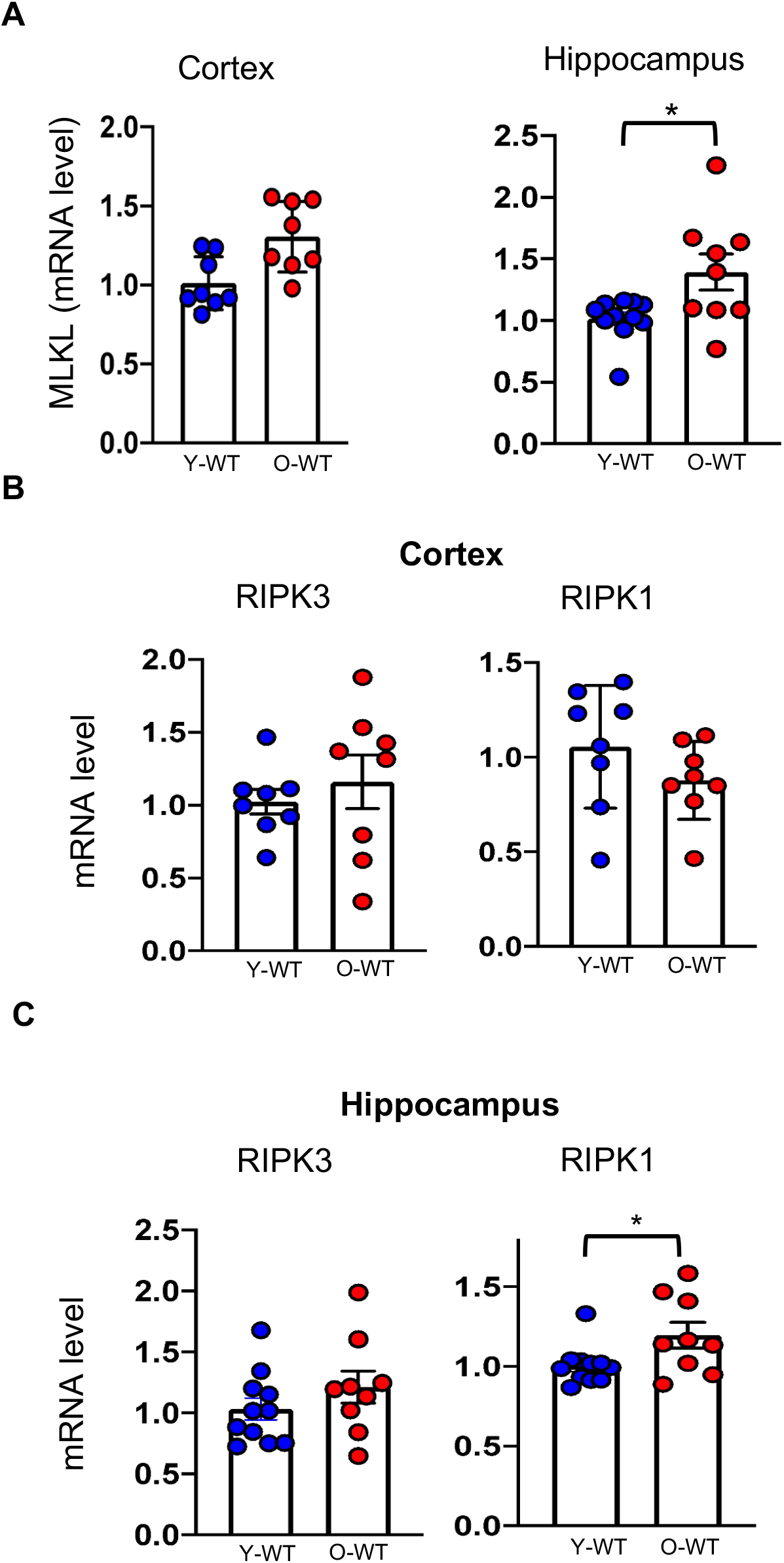
Transcript levels of MLKL in the cortex (left panel) and hippocampus (right panel) of young WT (Y-WT, red) and old-WT (O-WT, blue) mice **(A)**. Transcript levels of RIPK3 and RIPK1 in the cortex **(B)** and hippocampus **(C)** of Y-WT and O-WT mice. Data were obtained from 8 to 10 mice per group and are expressed as the mean ± SEM. * p ≤ 0.05).

**Supplementary Figure 2:**
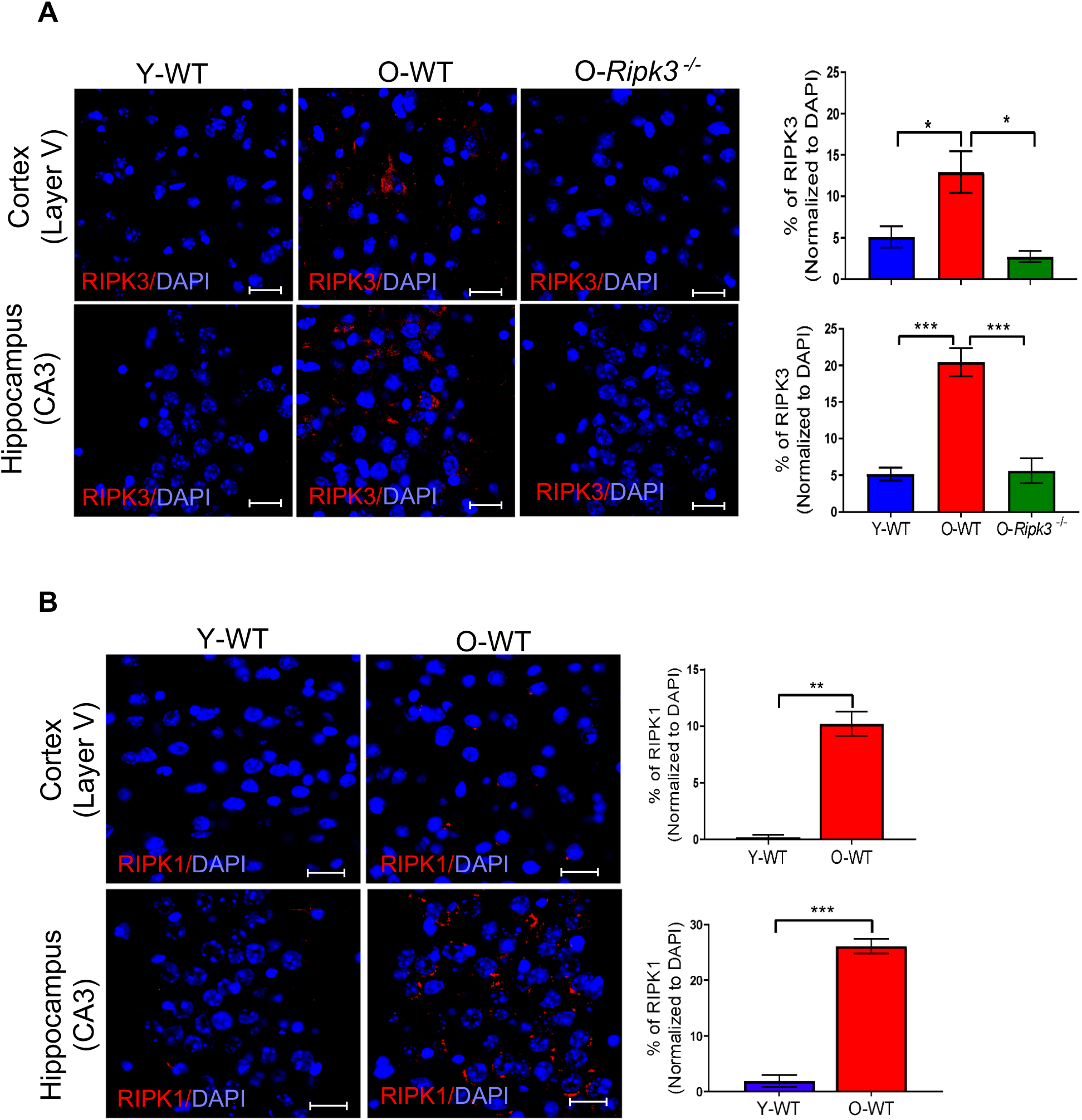
Confocal images showing expression of RIPK3 **(A)** and RIPK1 **(B)** in the layer V of the cortex and the CA3 region of the hippocampus of young WT (Y-WT), old WT (O-WT), and old *Ripk3*^*-/-*^ mice. RIPK3 and RIPK1 staining is in red and DAPI staining is in blue (left panel). Graphical representation of the percentage of RIPK3 or RIPK1 normalized to DAPI for Y-WT (blue bar), O-WT (red bar), and old *Mlkl*^*-/-*^ mice (green bar) are shown in the right panel. Data were obtained from 5-7 mice per group and are expressed as the mean ± SEM. * p ≤ 0.05, **p≤ 0.01, ***p≤ 0.001. Scale bar: 20 μm.

**Supplementary Figure 3.**
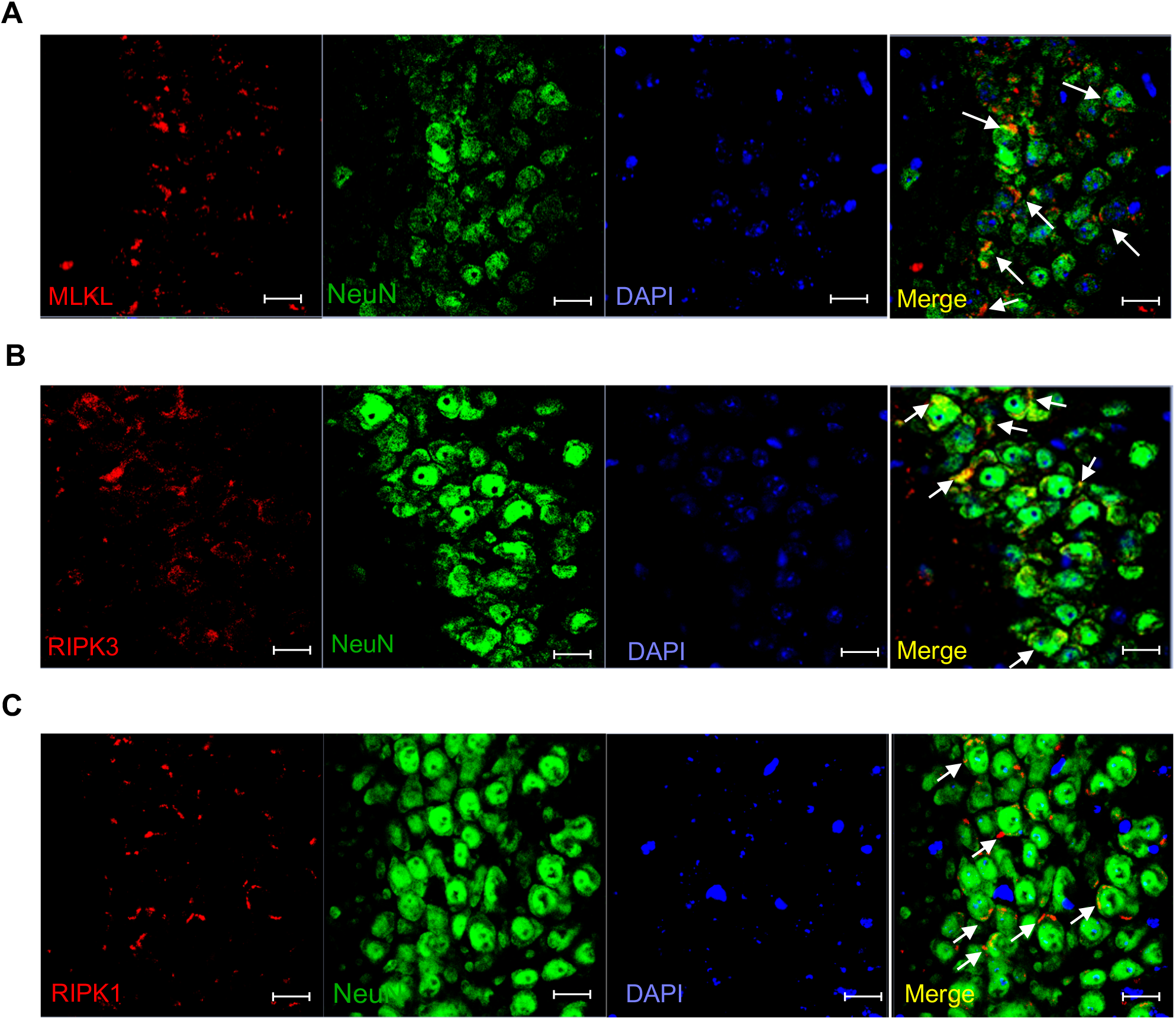
Confocal images showing expression of MLKL **(A)**, RIPK3 **(B)** and RIPK1**(C)** in neurons at the CA3 region of the hippocampus. Double immunofluorescence staining and confocal micrographs are shown for neurons (NeuN, green), MLKL or RIPK3 or RIPK1 (red) and DAPI (blue). White arrows indicate the co-localization of MLKL or RIPK3 or RIPK1 with NeuN. Scale bar: 20 μm.

